# Identification of antigenic epitopes on the F and G glycoproteins of bovine respiratory syncytial virus and *in vitro* assessment of their synthetic peptide vaccine potential

**DOI:** 10.1101/2021.06.25.449873

**Authors:** J. Lemon, A. Douglas, U. Power, MJ. McMenamy

## Abstract

Globally, bovine respiratory disease (BRD) remains the principal reason for mortality of calves over one month of age despite the availability of various vaccines on the UK market. Bovine respiratory syncytial virus (BRSV) was first discovered in the 1970s and is now considered a principal pathogen implicated in the disease complex. Outbreaks occur annually and re-infections are common even in the presence of maternal antibodies. Difficulties have arisen from using both live-attenuated and inactivated vaccines and recent efforts have focused on the development of sub-unit vaccines that are suitable for use in neonatal calves with maternally-derived circulating antibodies. This study was undertaken to identify antigenic epitopes on two of the surface glycoproteins of BRSV, the fusion (F) and attachment (G) proteins, the major surface viral antigens, for inclusion into a novel subunit peptide vaccine. Sequencing and antigenicity prediction of the F and G genes of BRSV revealed 21 areas of potential antigenicity; of which genuine peptide/antisera binding occurred with 4 peptides. Identification of the antigenic components of a vaccine is an important first step in the development of novel BRSV vaccines and this data, therefore, provides the basis for the generation of such vaccines.

## Introduction

Bovine respiratory syncytial virus (BRSV) is a member of the *Orthopneumoviridae* family and is a major respiratory pathogen of cattle, with the young and immunocompromised particularly vulnerable to infection. Enhanced pulmonary disease following infection was observed in calves (1) immunised using a formalin-inactivated alum-precipitated BRSV (FI-BRSV) which led to alternative antigens being investigated for use in BRSV vaccines.

The two major glycoproteins of BRSV are the major attachment protein (G) and the fusion protein (F), both of which are targets for virus neutralising antibodies (VNA). The G protein is heavily involved with the modulation of the host immune response, through mechanisms such as glycosylation, receptor mimicry and antibody interference (2-4) while the F protein is known to confer cross-protection between strains due to its high degree of sequence conservation. The F2 subunit has also been suggested as being responsible for the species-specificity witnessed with RSV infection (5).

Various antigenic epitopes within either protein have been identified. Taylor *et al* (1997) reported reduced pneumonic lung lesions and the efficient induction of bovine CD8+ T cells after immunisation with a recombinant vaccinia virus (rVV) expressing the F protein (6). In addition, Fogg *et al* demonstrated T-lymphocyte proliferation using radioactively-labelled PBMCs which had been harvested from previously vaccinated calves and subsequently incubated with peptides (Fogg, Parsons, Thomas, & Taylor, 2001). In an earlier study calves were immunised with a peptide spanning residues 174-187 from the G protein of BRSV 375 strain and, although no statistical difference was noted between viral titres in lung homogenates and washings, no clinical signs of respiratory disease were observed and reduced lung consolidation comparative to infected animals was observed at pathology (Bastien et al., 1997).

Epitope mapping has also been used to define immunopathological residues to avoid; information which could complement epitopes presented for use in a subunit vaccine. Enhanced lung eosinophilia has been noted after vaccination using FI-RSV antigen and subsequent infection. Using two pGEM4-derived plasmids encoding the G proteins of frameshift mutants, Sparer et al identified residues G193-203, located in the carboxyl terminal of RSV G, as responsible for this. In the same study, mice which had been primed intranasally with recombinant Vaccinia virus incorporating the same residues displayed similar weight loss to those vaccinated with wild-type G, both of which groups demonstrated statistically greater weight loss than negative control vaccines (7).

The first reported example of a veterinary peptide vaccine providing protection in the natural host was developed against canine parvovirus (8). Since then peptide vaccines have been assessed for their efficacy in various species against many other veterinary viral pathogens, such as foot and mouth disease virus (FMDV) in sheep (9), classical swine fever virus in pigs (CSFV) (10) or bovine papillomavirus-4 in calves (11). Recently, there has been a resurgence of interest in options for synthetic vaccines, including investigating peptide technology, related to the production of safe, efficacious Covid-19 coronavirus vaccines (12, 13).

As a vaccine must establish an antibody response, it is essential to identify B-cell epitopes for inclusion in novel sub-unit vaccines. This details the selection and *in vitro* screening of synthetic peptides derived from the BRSV F and G proteins for antigenicity against a panel of bovine sera positive for anti-BRSV antibodies.

## Materials and methods

### Virus Propagation

BRSV strain 274 used in this study was kindly donated by Prof. M. Elvander, SVA, Sweden. It was isolated from an adult cow showing signs of severe respiratory distress during a Swedish outbreak in 1997. A virus pool was grown in foetal calf lung cells (FCL), a semi-continuous cell line obtained in AFBI (VSD, NI). GMEM-BHK 21 1X containing 1% L-glutamine (200mM) minus Tryptose Phosphate Broth was supplemented with 10% heat-inactivated Foetal Bovine Serum and 0.1% Gentamicin (50mg/mL). Cells were cultured in 75 cm2 flasks at 37°C in 5 % CO2 in 18 mL growth medium and on reaching confluency, cells were passaged 1 in 3 every 7 days. Once they reached around 70 % confluency (∼4 d after passage) the growth medium was removed and 1 mL BRSV stock was adsorbed to the monolayer for 1.5 h at 37°C / CO2. A control flask was treated with PBS only. The virus inoculum was not removed and after incubation, 19 mL of maintenance medium was added to the flask. Cells were inspected daily and harvested upon observation of extensive CPE (∼6 d post-infection) and stored in cryovials at -80 °C.

### Viral RNA extraction

RNA from BRSV stocks were extracted using a QIAamp Viral RNA mini kit (Qiagen, UK) as per manufacturer’s instructions.

### Obtaining BRSV F and G gene sequence

Endpoint RT-PCR was performed on extracted viral RNA using primers designed in-house amplifying areas outside the 3’ and 5’ ends of the F and G genes. These are detailed in Appendix 1. For RT-PCR, primers were used at a concentration of 10 μM and primer sets G1/G2, F1/F3, F2/F4 were used. All reactions were set up in triplicate using a One-Step RT-PCR kit (Qiagen, UK) as per the following run conditions: 30 mins @ 50°C; 15 mins @ 95°C followed by 30 cycles of 30 sec @ 94°C; 30 sec @ 51°C; 2 mins @ 72°C followed by 10 mins @ 72°C. Amplicons were electrophoresed on a 1 % agarose gel at 120 V for 60 min. Amplicons of appropriate size were excised and purified using an Isolate II PCR and Gel kit (Bioline, UK). The purified amplicons were included in cycle sequence reactions, prepared using primers as detailed before, only at the concentration of 3.2pmol and a Big Dye Terminator kit (Life Technologies, California), as per manufacturer’s instructions. Sequencing was performed on a 3730 XL genetic analyser (Thermo Fisher, UK) and analysis performed using Geneious v10.0.

### B cell epitope prediction

Translated amino acid sequence was determined from the DNA sequence, using Geneious R 9.1.3. (Biomatters, NZ). To predict antigenic epitopes, this data was entered into the Immunomedicine online prediction software (http://imed.med.ucm.es/Tools/antigenic.pl IMED, Spain) the results of which were cross-referenced with current literature and alternative antigenicity prediction software (National Centre for Biotechnology Information; NCBI). This software produces *in silico* models of hydrophobic and hydrophilic areas of sequence, thus identifying potential areas of antigenicity.

### Peptide synthesis

Using current literature and prediction software, 21 peptides were chosen for further analysis in this study. A single amino acid residue substitution between the sequence for peptides 1, 2 and 3 allowed us include a sequences from 2 Danish strains of BRSV (strain 9402022, strain 9416116) providing some heterogeneity for the study. Peptides were synthesized commercially by Mimotopes, Australis and were synthesised at >70 % purity. All peptides, except peptide 4, were labelled with biotin at the N terminus and capped with an amide group (NH2) at the C terminus. All peptides contained a 4 mer serine-glycine-serine-glycine (SGSG) spacer following the biotin tag at the N terminus to reduce the potential for steric hindrance, thus allowing the necessary epitope for antibody binding to be accessible. As peptide 4 was the starting sequence of the G protein, synthesized without a biotin tag, it was hypothesised that a free amine group at the N terminus would more closely represent an N-terminal native epitope and thus might be more beneficial for antibody binding than a biotin tag. The selected peptides are detailed in Appendix 2.

### Selection of bovine sera

A panel of bovine sera were kindly supplied by the Agri-Food and Biosciences Institute Diagnostic Surveillance and Investigation Branch in Veterinary Sciences Division (DSIB, VSD, AFBI, Belfast). These were obtained from diagnostic submissions, as part of the NI passive disease surveillance program for the Department of Agriculture, Environment and Rural Affairs (DAERA). Sera were screened using commercial indirect ELISA kits (Svanova, Sweden) for antibodies against bovine respiratory syncytial virus and bovine parainfluenza virus type 3.

### Optimisation of ELISA protocol prior to screening

Optimisation work was performed to determine an appropriate plate type and peptide coating concentration. All dilutions were prepared in PBS 1X pH 7.4.

- Selection of plate type: The following plates were used: Nunc Maxisorp C bottom, Pierce Streptavidin-coated, Microlon 200 U bottom and Immulon 1B plates. These ranged from ranged from high to medium affinity and a variety of coating methods were assessed: 1/ 100 μL 0.025 % (v/v) glutaraldehyde (Sigma, UK) which was incubated at 37 °C with shaking for 1h. This was removed and 50 μL peptide was added along with 10 μL water to ensure consistency in volume with the other test methods 2/ 10 μL 0.025 % (v/v) glutaraldehyde was added to wells containing 50 μL peptide. 3/ 50 μL peptide was added directly to the plate, with the addition of 10 μL water to ensure consistency in volumes with the other test methods. All peptides were coated at 5 μg/mL or 1 μg/mL in and left to incubate at +4°C overnight. The following morning, the ELISA protocol, as described in Appendix 3, was performed.
- Coating optimisation: The lowest molecular weight peptide was coated at 10, 5, 1 and 0.1 μg/mL across the chosen plate, which was then sealed and left overnight at +4°C. The following morning a washing step was performed and biotinylated HRP, containing 1 % bovine serum albumin (v/v) was dispensed at 50 μL/well in decreasing concentrations of 1:80,000, 1:160,000, 1:320,000 to 1:640,000 down the plate. Blocking buffer was removed to reduce any associated non-specific binding and the ELISA protocol, detailed in Appendix 3, was followed from step 6.

### *In vitro* antigenicity screening using bovine sera

All 21 peptides were coated at 5 μg/mL and screened using a pool of BRSV antibody-positive and bovine negative sera, using the ELISA protocol detailed in Appendix 3.

### Repeat testing of peptides using anti-BRSV antibody negative sera

All peptides were tested using bovine sera which did not contain antibodies against BRSV to ensure any reactivity observed during initial screening was specific for BRSV and not as a result of non-specific binding.

### Acceptability criteria

At initial analysis using antibody positive sera, any peptide which produced optical densities (OD) less than or equal to twice the negative control value was eliminated, as reactivity was considered too low to inform any further investigations. Corrected averages were then obtained by removing the negative control value, using both antibody negative and antibody positive sera from the respective optical density results, to negate background optical density. A ratio was calculated between the optical density obtained for an individual peptide using antibody positive sera and antibody negative sera and peptides were considered as displaying genuine reactivity if this value was >2.

## Results

### G and F gene sequencing and translated protein sequence

The sequences of the BRSV G and F genes were obtained using a 3730 XL genetic analyser (Thermo Fisher, UK) and amino acid sequences of the respective G and F proteins predicted using Geneious v 9.1.3 (Biomatters, NZ). The nucleotide sequence of the BRSV G and F genes and the predicted amino acid sequence of the BRSV G and F proteins that were entered into selected online antigenicity prediction software tools have not been published before and are found in Appendix 4. Phylogenetic trees are shown in Figures 1 and 2, which demonstrate the genetic similarity of BRSV F and G genes from the reference strain 274 used during this study in comparison to the F and G genes of other strains of BRSV.

**Figure 1:**
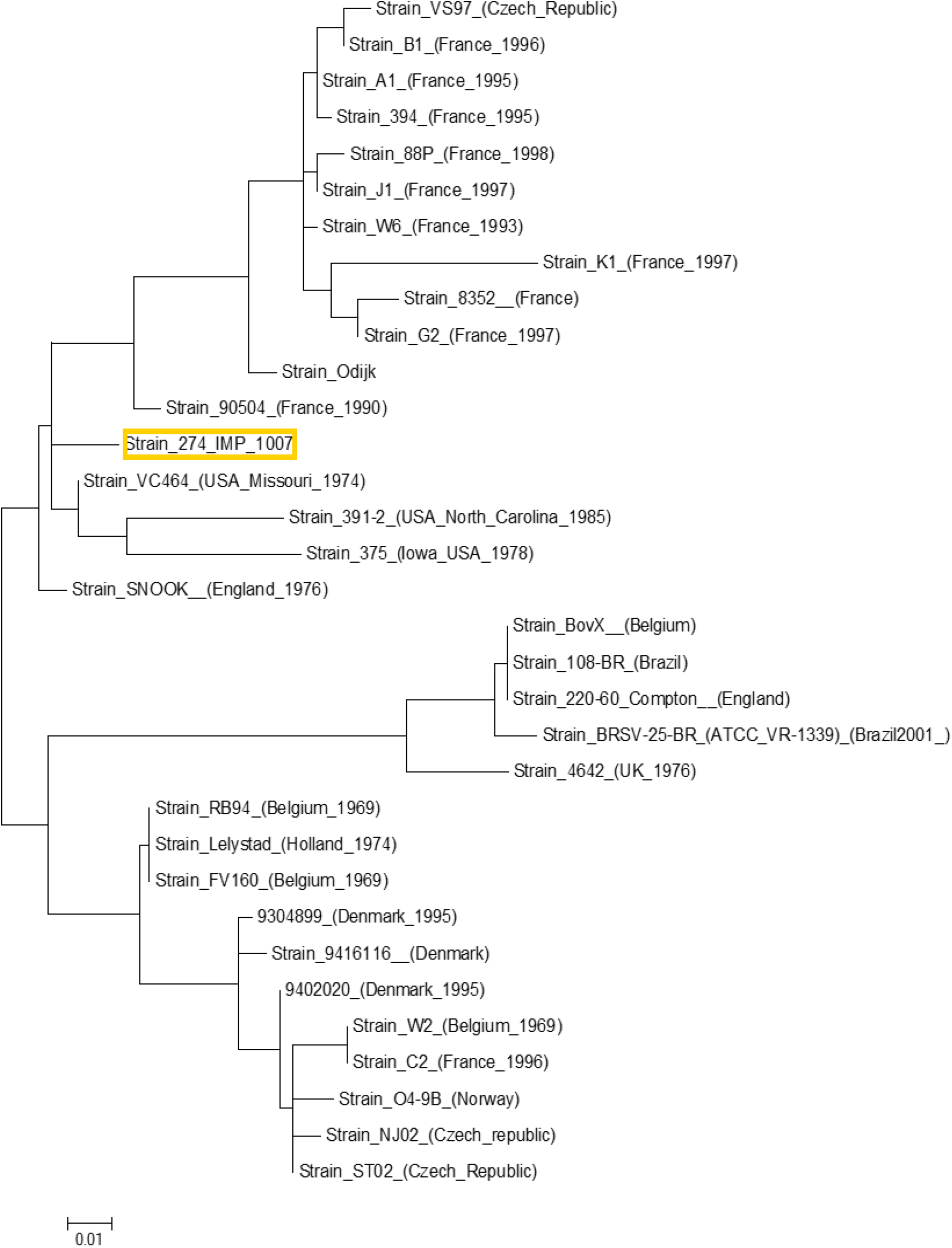
Phylogenetic tree (cladogram) of published G gene sequences, obtained from NCBI demonstrating how the reference strain 274 (IMP 1007; outlined in yellow) from this study relates to other strains of BRSV. Branch lengths are proportionally representative of the amount of evolutionary change.

**Figure 2:**
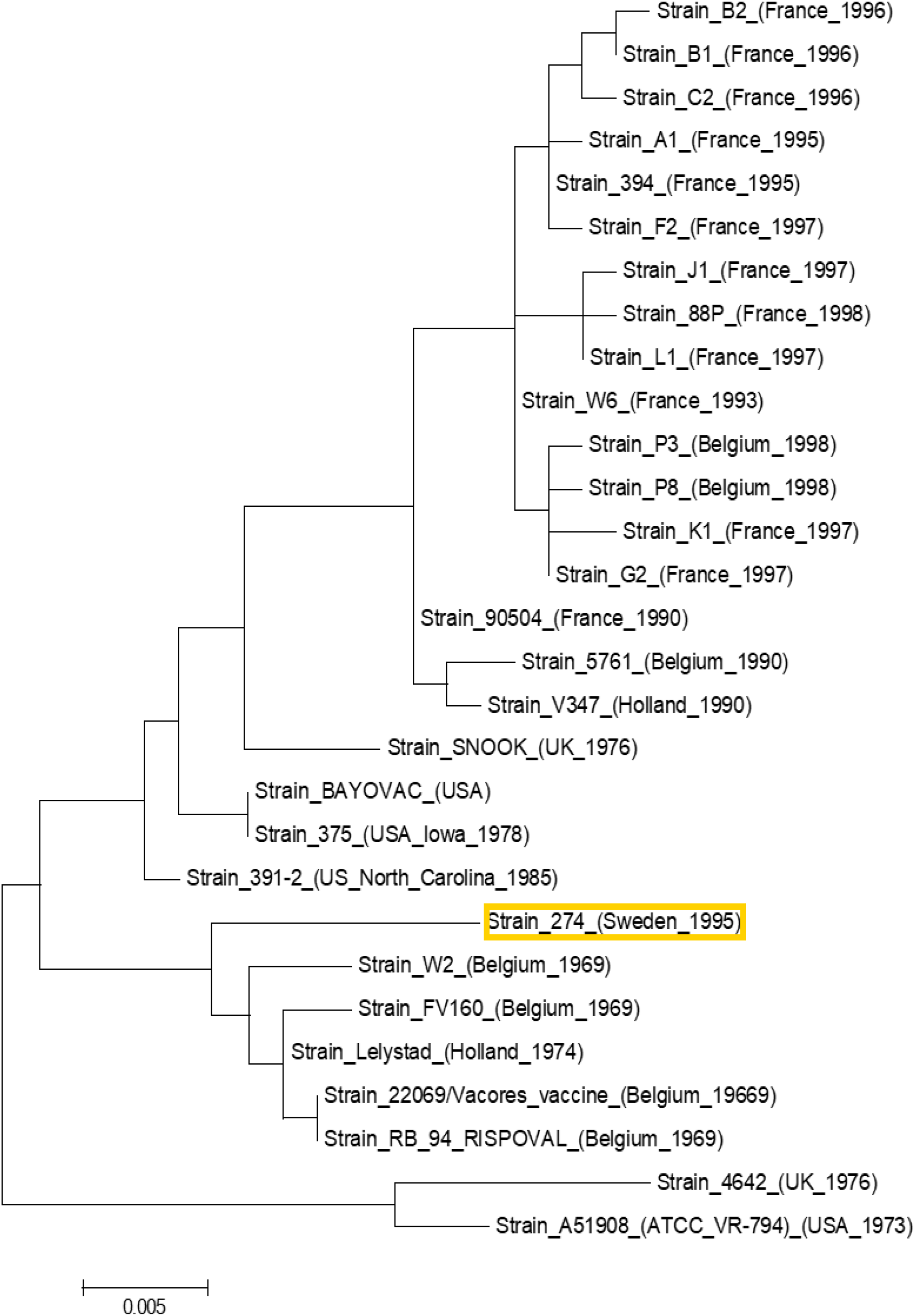
Phylogenetic tree (cladogram) of published F gene sequences, obtained from NCBI demonstrating how the reference strain 274 (Sweden 1995; outlined in yellow) from this study relates to other strains of BRSV. Branch lengths are proportionally representative of the amount of evolutionary change.

### Peptide selection

Predicted antigenic peptides were obtained using the online antigenicity prediction software (http://imed.med.ucm.es/Tools/antigenic.pl) as previously described. Results obtained using this software were then cross-referenced using a different online prediction tool (NCBI) and with sources taken from the literature. The IMED-generated antigenicity charts are shown in Figures 3 and 4 for the BRSV G and F proteins, respectively. Peptides with an antigenic propensity (i.e. potentially antigenic) ≥1.0 spanning >10 amino acids were selected. The chosen sequences corresponded to the peptides defined in Appendix 2, which lists the BRSV protein containing the sequence, the residue locations on the protein, the predicted amino acid sequence of the chosen peptide and the theoretical molecular weight (in Daltons, Da). All peptides were given a unique in-house identification number.

**Figure 3:**
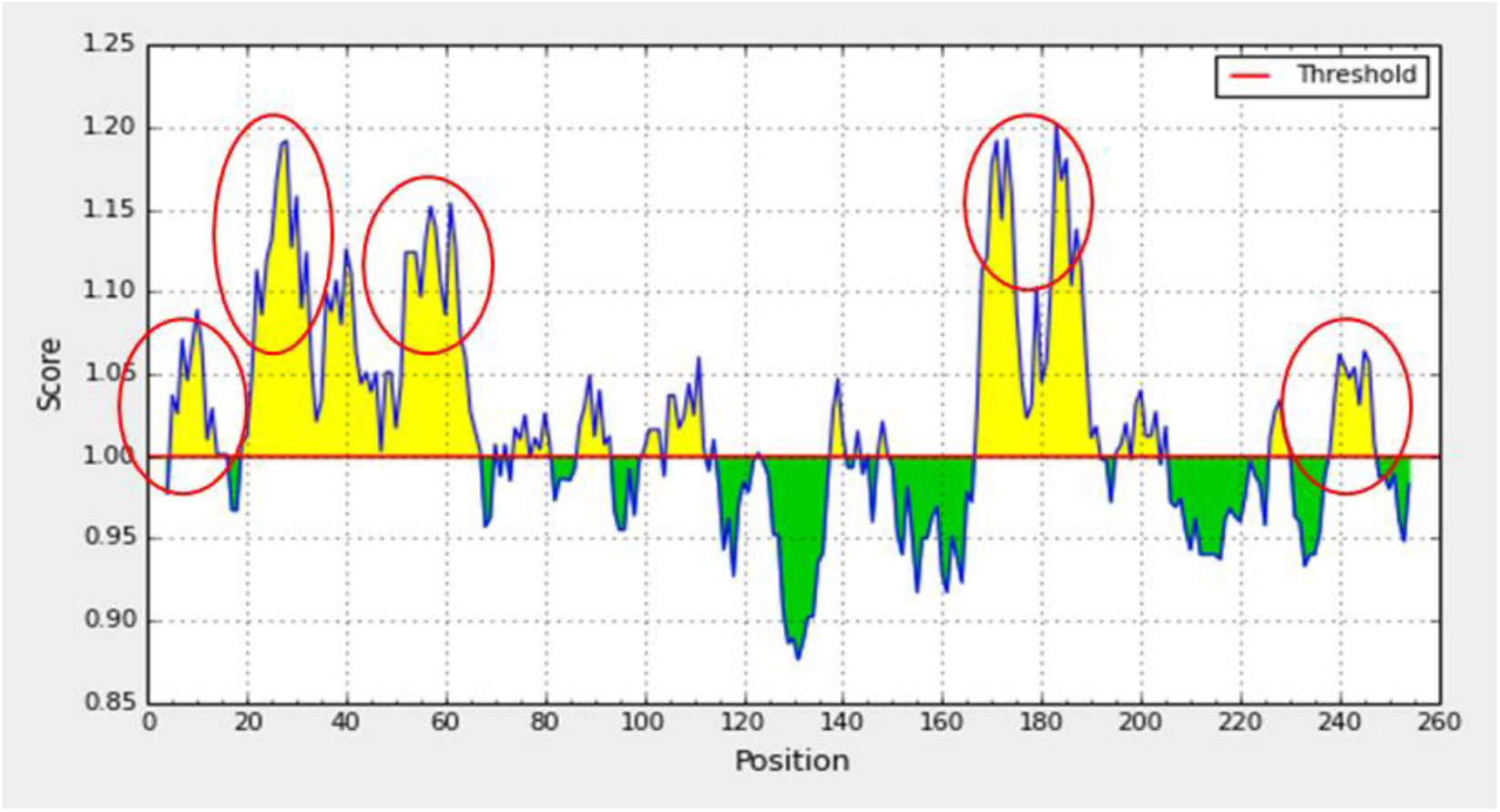
Predicted antigenicity chart for BRSV G protein. Red circles indicate the areas where selected peptides are located. Yellow bars indicate areas above the threshold (solid red line) with a positive score of 1.00 or above. The position on the primary sequence is noted along the X axis of the graph. Anything in solid green was disregarded.

**Figure 4:**
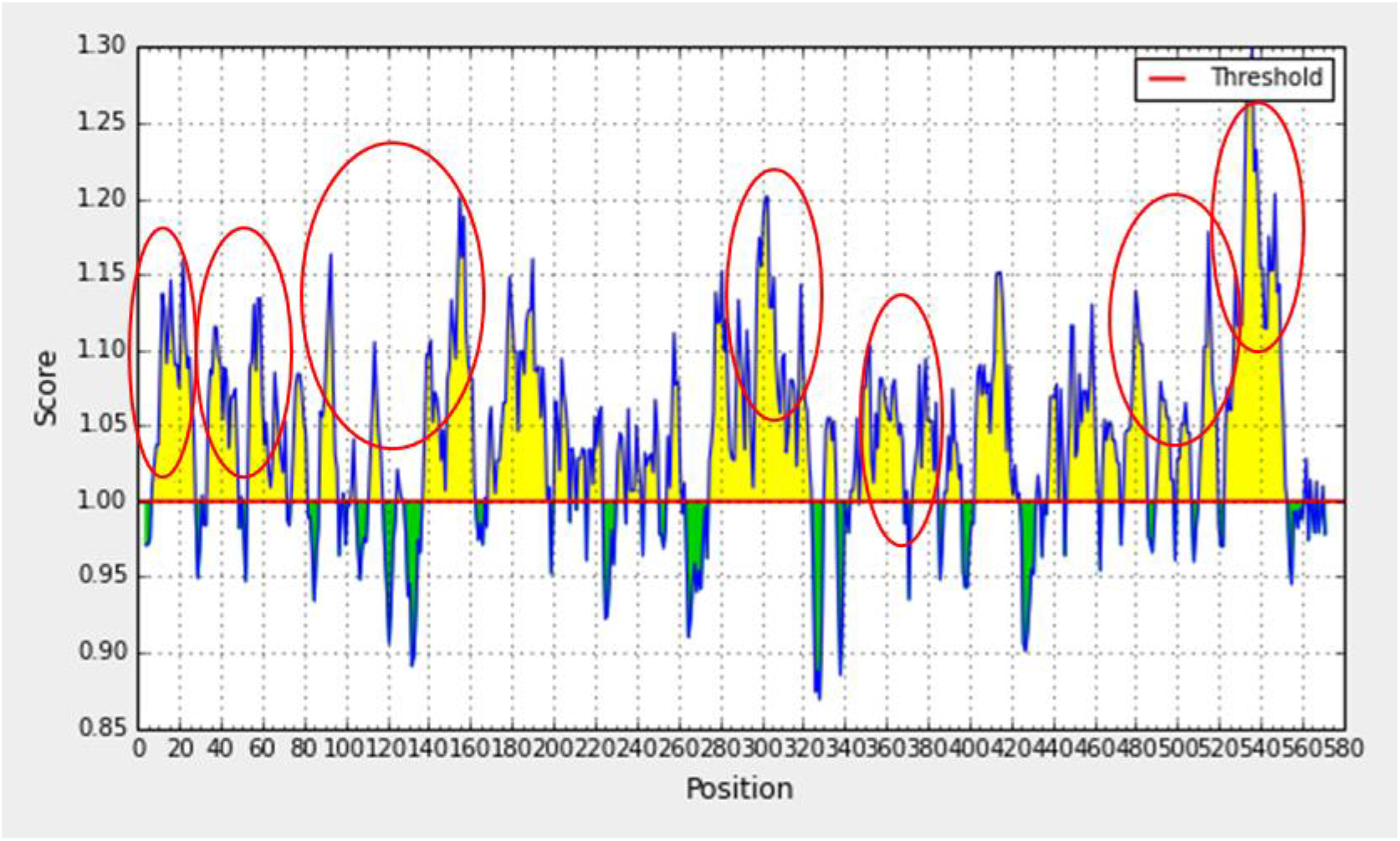
Predicted antigenicity chart for BRSV F protein. Red circles indicate the areas where selected peptides are based. Yellow bars indicate areas above the threshold (solid red line) with a positive score of 1.00 or above. The position on the primary sequence is noted along the X axis of the graph. Anything in solid green was disregarded.

### Optimisation of ELISA protocol prior to screening

Optimisation work was performed to determine an appropriate plate type and peptide coating concentration. High levels of non-specific binding, regardless of peptide coating concentration or method, was observed when using Microlon or Nunc plates. Immulon 1B plates also failed to produce acceptable results and O.D. values consistently >1 suggested limited binding, regardless of the test method. Using the Pierce Streptavidin-coated high binding capacity (HBC) clear 96-well plate, coating the plate at 5 μg/mL resulted in the greatest difference between the peptide-coated well and the negative PBS-coated OD450 reading, thus allowing a greater capacity to differentiate between the test peptides (data not shown) For this reason, these plates were chosen for all subsequent experiments. A minimum coating concentration was determined by performing chequerboard analysis using the lowest molecular weight peptide in varying concentrations, ranging from 10-0.5 μg/mL. A coating concentration of 5 μg/mL was chosen (data not shown). Results suggested that coating at less than 5mg/mL produced a limited O.D. range for future experiments while little difference was noted between O.D. values when coating was performed at 10 mg/mL or 5 mg/mL.

### *In vitro* antigenicity screening

Initial screening of all peptides by ELISA indicated that 13 of the selected 21 peptides displayed reactivity when tested with BRSV antibody positive sera, as they were above the cut-off value defined previously. Seven peptides were eliminated because the optical densities were ≤2x the negative control value (0.22). Figure 5 displays the optical densities obtained when peptides were screened with BRSV antibody-positive sera. Subsequently all 21 peptides were tested with bovine sera which had previously been determined as not containing antibodies against BRSV. When the ratio was calculated between the result obtained using negative and positive sera for an individual peptide, 4 peptides (#8, 9, 10 and 19) were above the previously defined cut-off ratio. Several peptides within the G protein sequence were determined as having very high ratios and 1 peptide within the F protein sequence (#19) had a ratio just above the cut off value of 2 and requires further investigation. Additionally, this negates the reactivity evident when testing with antibody positive sera for peptides #1-3, 6 and 12-17, which had originally met acceptability criteria. The ratio is displayed as a line graph in Figure 6 and the 4 peptides identified as antigenic in this study are detailed in Table 1. Figures 7-10 show several alignments of the 4 selected peptides to demonstrate if they are conserved amongst BRSV strains

**Figure 5:**
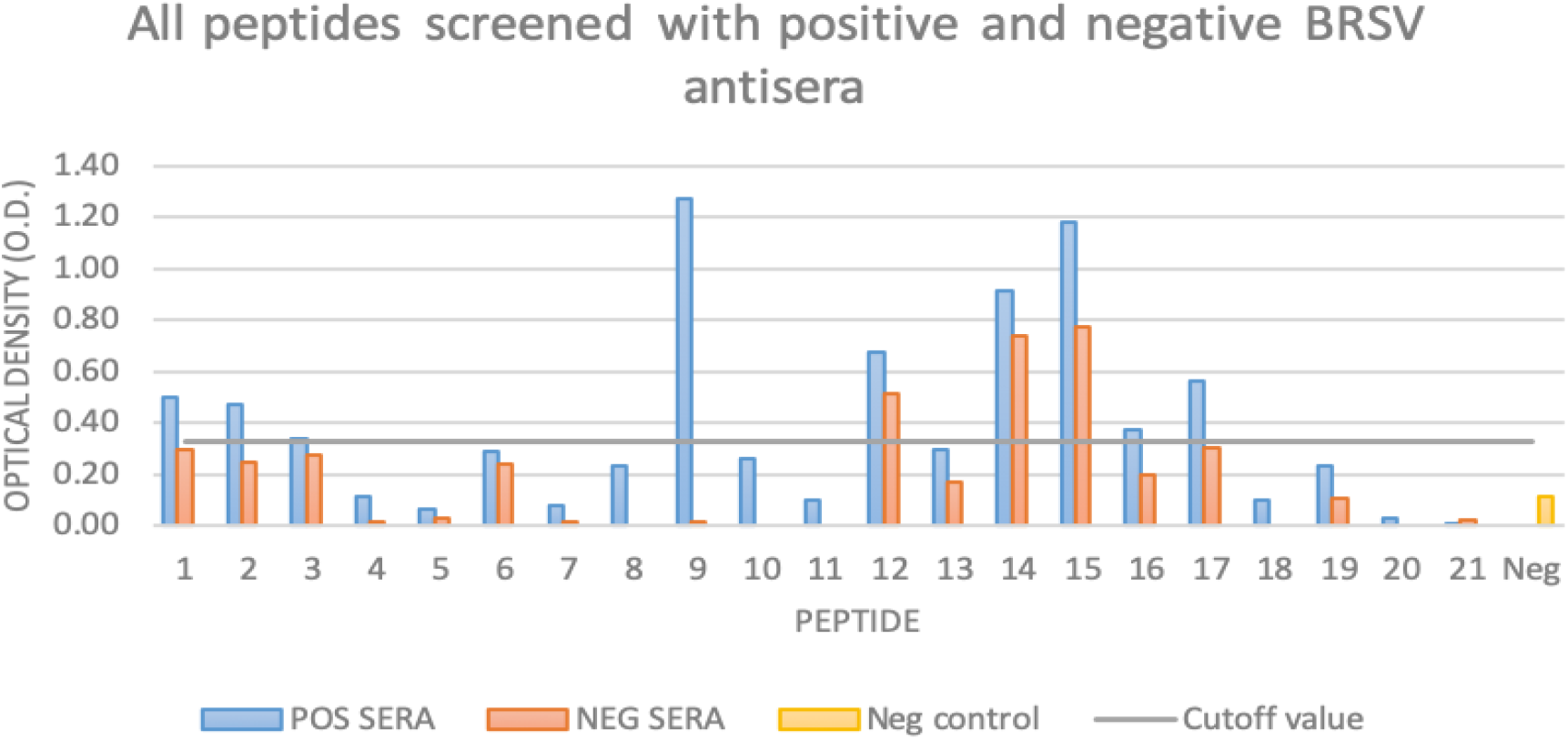
Bar chart of all peptides when screened with BRSV antibody positive sera (blue bars) and BRSV antibody negative sera (orange bars). The solid grey line line represents the cut-off point, below which optical densities were deemed too low to be considered reactive (negative control x2; ≤0.22).

**Figure 6:**
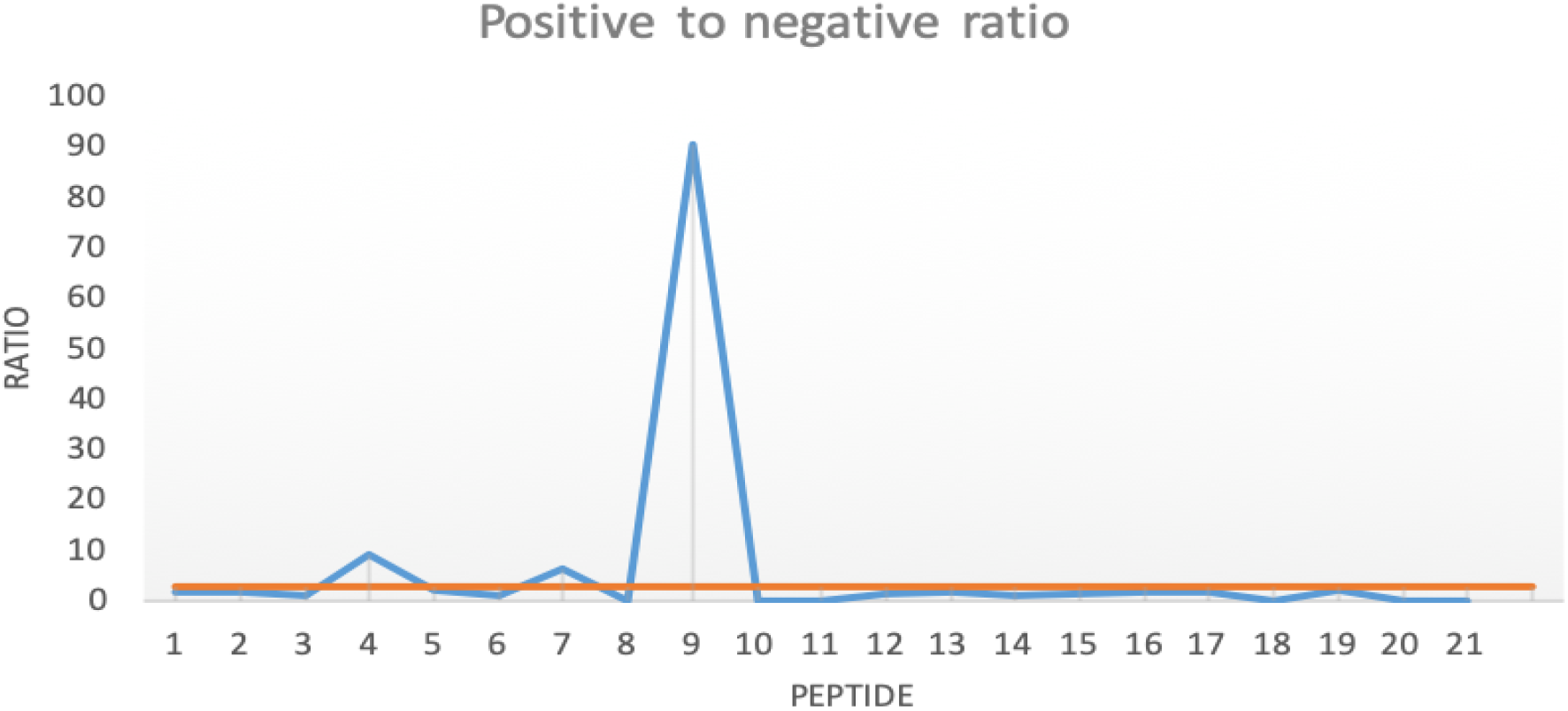
Line graph demonstrating the ratio between the optical densities obtained from testing the selected peptides with BRSV antibody-positive and negative sera. The blue line represents the ratio while the orange line represents the cut-off value (≥2).

**Table 1:**
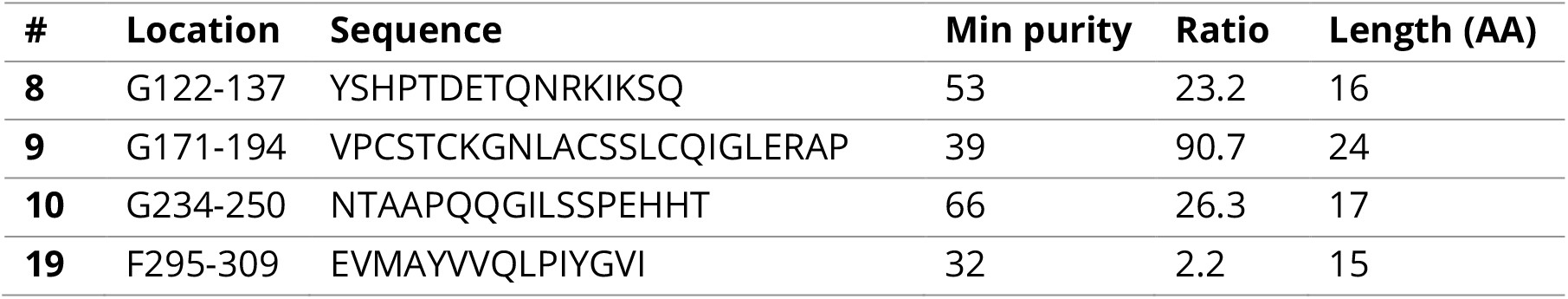
The 4 peptides which displayed genuine reactivity, as defined by the acceptability criteria, are listed, along with the minimum purity, ratio value and amino acid length. All peptides had a biotin tag and an SGSG spacer prior to the amino acid sequence at the N terminus

**Figure 7.**
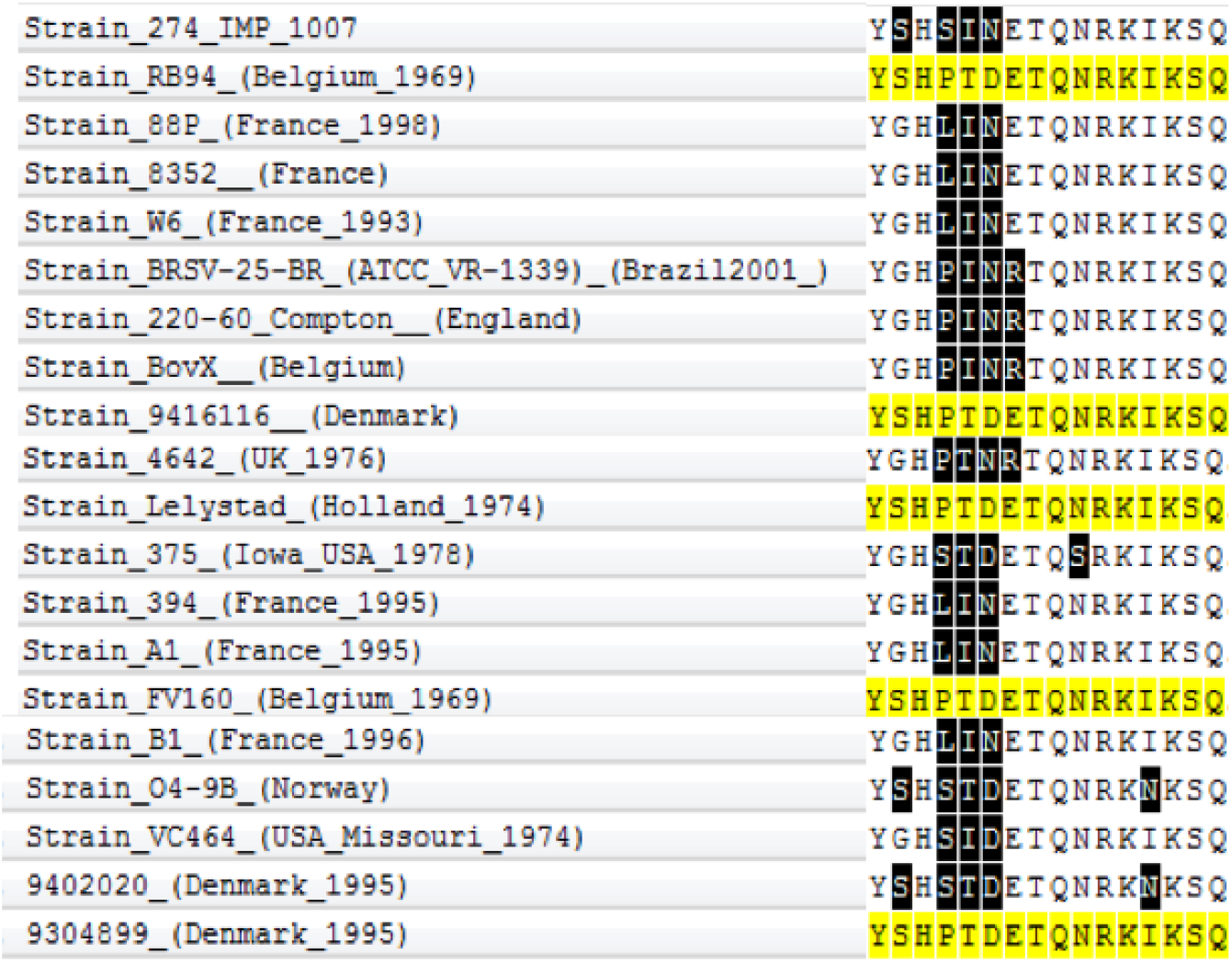
Twenty BRSV strains were aligned at residues G122-137. Sequence identities ranged from 69 - 100 % identity to the antigenic peptide which was chosen from the Rispoval vaccine strain (RB94, Belgium). 100 % homology is demonstrated with 4 other strains, indicating that this is a conserved epitope within the G protein.

**Figure 8.**
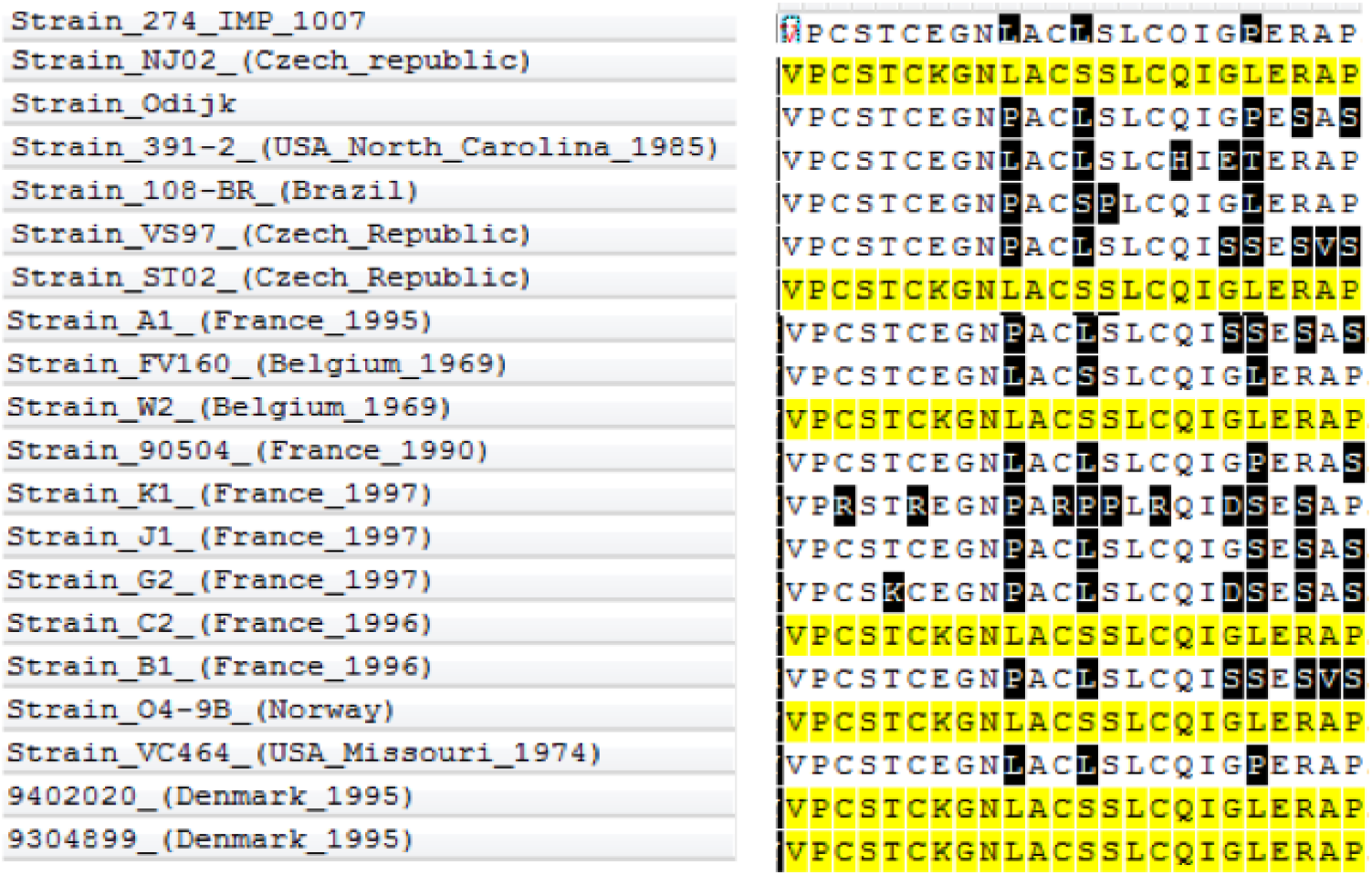
Twenty BRSV strains were aligned at residues G171-194. Sequence identities ranged from 58 – 100 % identity to the antigenic peptide which was chosen from the Czech_republic strain NJ02. 100 % sequence identity is demonstrated with 6 other strains (highlighted) indicating that this is a conserved epitope within the G protein.

**Figure 9.**
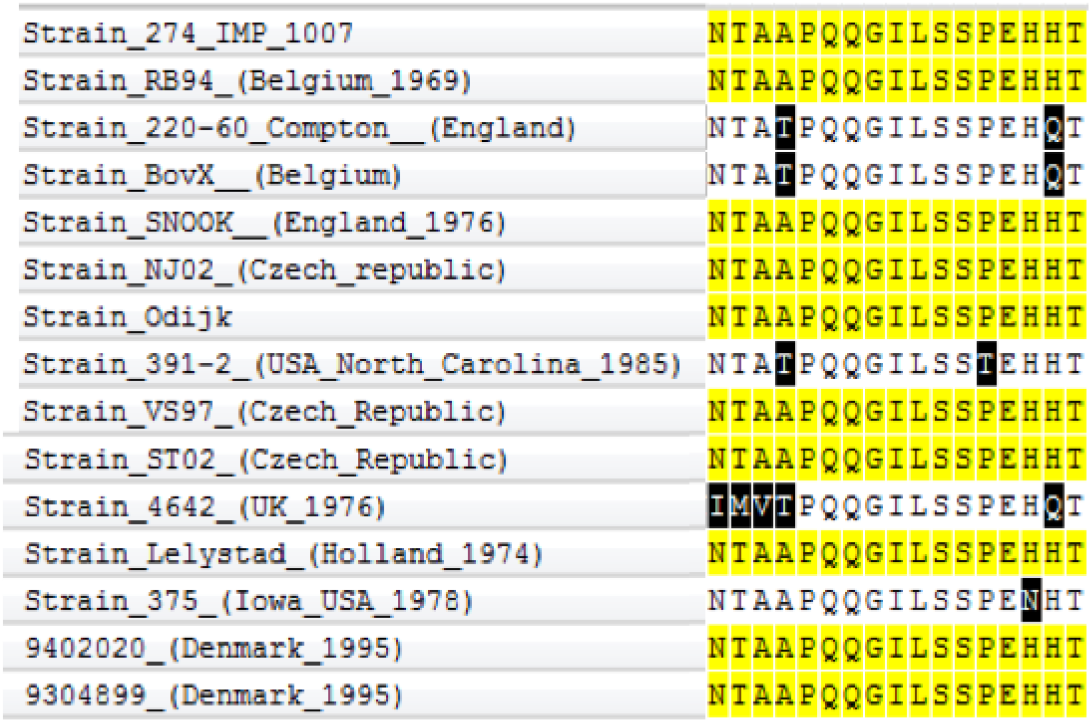
Fifteen BRSV strains were aligned at residues G234-250. Sequence identities ranged from 87 - 100 % identity to the antigenic peptide chosen from the reference strain (274_IMP 1007). 100 % sequence identity is demonstrated with 9 other strains (highlighted) indicating that this is a highly homologous epitope within the G protein.

**Figure 10.**
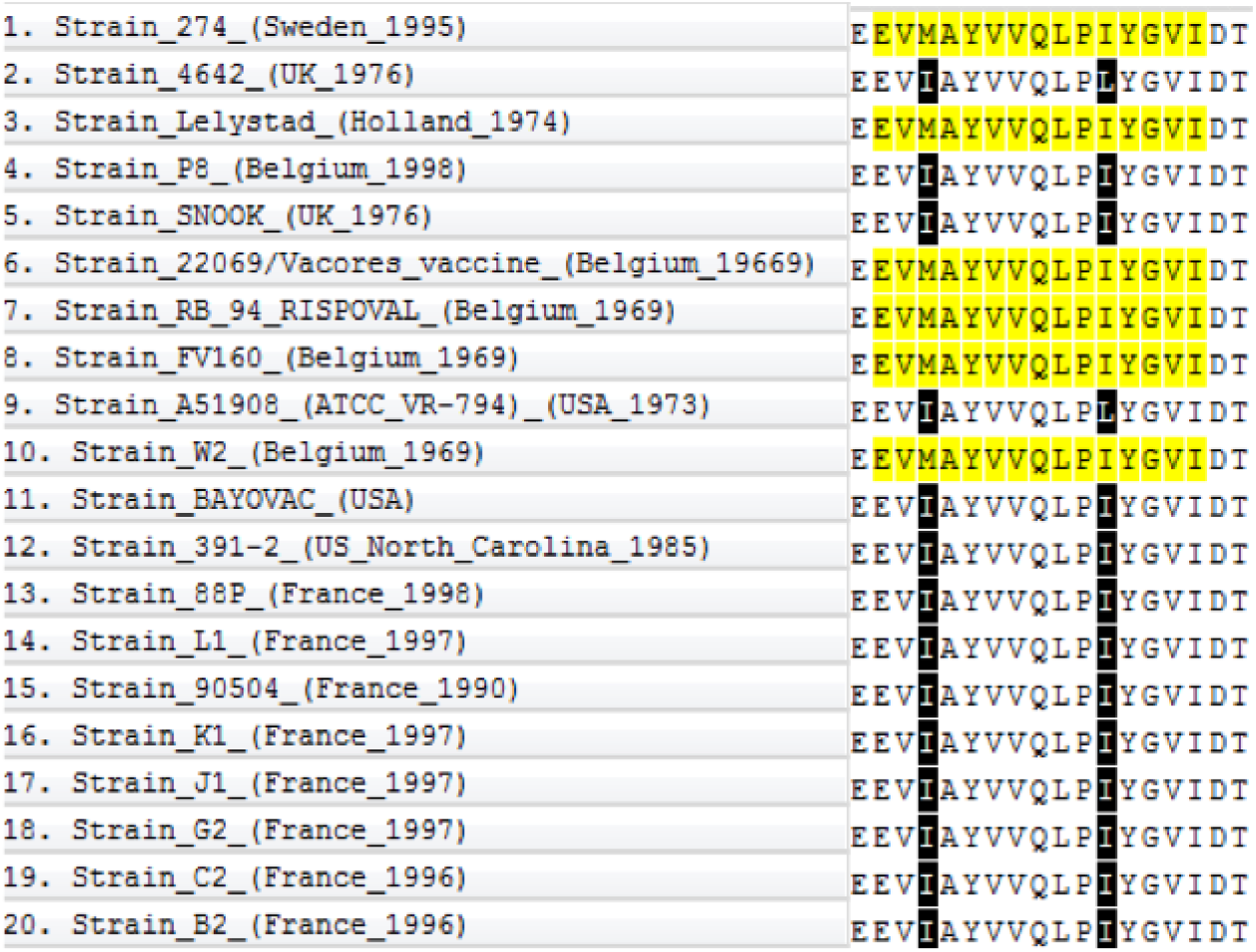
Twenty BRSV strains were aligned at residues F295-309. Sequence identities ranged from 87 – 100 % identity to the antigenic peptide chosen from the reference strain (274_IMP 1007). 100 % sequence identity is demonstrated with 5 other strains (highlighted) indicating that this is a highly conserved epitope within the F protein.

## Conclusion

The development of reverse genetic systems and recombinant DNA technologies have provided scientists the means to develop an unprecedented level of knowledge about the function and structure of RSV proteins. Consequently, it is now feasible to dissect the immune responses directed against viral proteins down to the amino acid level. Domains corresponding with the antigenic regions or epitopes of the proteins may be reproduced by synthetic chemistry, thereby providing a rational approach to vaccine development.

As new virus strains evolve through genetic mutations, with the continuous evolution of BRSV F and more specifically G sequences that has been previously reported (14), current conventional vaccines may become less efficacious. However, because of the ease with which we can sequence viral genomes, alternative epitopes can be easily and quickly identified. Of particular relevance to the veterinary field is the potential use of peptides in DIVA vaccines, also known as marker vaccines; vaccines intended to differentiate between vaccinated and infected animals (15). These have shown much promise with Foot and Mouth Disease (FMD), Classical Swine Fever (CSF) and Avian Influenza (AI) and are particularly valuable to support trade in countries with disease-freedom status (16).

Recent trends shifting the focus from the use of live-attenuated or inactivated whole virus as vaccine antigens mean sub-unit vaccines against BRSV, including those based on peptides, are gaining prominence. Peptide vaccines have many advantages in terms of safety over current vaccines. There is no infectious material involved and thus the same risk posed by live or inactivated vaccines, potentially resulting from the maintenance of virulence due to insufficient attenuation or inactivation, respectively, is eliminated. Furthermore, biochemical modifications, such as the addition of a lipid, carbohydrate or phosphate groups, can be introduced in a synthetic manner to improve immunogenicity, stability and/or solubility. From a practical perspective, peptide synthesis on a large-scale is economical and ensures less batch to batch variation, thereby facilitating completion of quality controls of vaccine efficacy (15). Several antigenic epitopes of BRSV have also been discovered and candidate peptide vaccines against BRSV have shown promise thus it was hypothesised that a peptide approach would provide a viable option for a BRSV vaccine. This study looked at the *in vitro* screening of a range of linear peptides in an attempt to define B-cell antigenic domains that bind antibodies naturally raised against the major surface glycoproteins of BRSV, G (attachment protein) and F (fusion protein) and is the first essential step needed to progress next generation vaccines from those which use the entire viral coding sequence.

As demonstrated in Appendix 4, 3 defined regions can be observed in the primary structure of the RSV G protein; the cytoplasmic tail (1-37 AA), the transmembrane region (38-65 AA) and the extracellular domain (66-257 AA). The protein is post-translationally glycosylated, which contributes to immune evasion, and within the extracellular domain there are two mucin-like regions flanking a highly conserved unglycosylated domain (17, 18) spanning G_163-189_. Many known antigenic epitopes are located in the mucin-like C domain or in the central conserved domain (17, 19, 20) including one of the peptides identified as antigenic during this study (G_171-194_) underlining the importance of the carboxyl terminus of RSV G in the immunogenicity of the protein.

A number of studies demonstrated the importance of epitopes located within the unglycosylated conserved domain in affording protective efficacy. Immunisation of mice with a peptide spanning residues 174-187 of the HRSV, subgroup A G protein conjugated to Keyhole Limpet Haemocyanin induced high titres of RSV-specific antibodies and protected mice from HRSV challenge (21). Later, Power *et al* demonstrated that a bacterially-expressed HRSV A G fragment (G_130-230_) fused to the albumin binding domain of Streptococcal protein G, denoted BBG2Na, provided protective and durable immune responses in mice for at least 46 weeks (22). The same group also employed this peptide with alternative carrier proteins (23, 24) or demonstrated the protective efficacy afforded to pups by maternal vaccination using BBG2Na (25).

Reactivity was observed between the BRSV antibody positive sera and peptides G_122-137_ and G_234-250_ during this study. These are located in the variable regions on either side of the central conserved domain and examples of the antigenicity of these specific residues are limited in the literature. However, recently a recombinant nanoparticle vaccine based on the entire ectodomain of BRSV strain 375 G protein was described (26). Reduced shedding, decreased viral loads and diminished pneumonic lung consolidation were reported, alongside increased BRSV-specific IgG serum titres, bovine dendritic cell activation and clonal expansion of CD4 T cells after immunisation of 2-3 week old calves (26). Whether or not the particular peptides identified in this current study are involved in the immunogenicity of this vaccine could only be determined by pepscan analysis using overlapping peptides which cover the entire ectodomain of the protein.

However, it should be noted that peptides G_122-134_ and G_234-250_, while antigenic, also contain potential acceptor sites for N-linked sugars, although the impact of glycosylation on vaccine antigen immunogenicity is not well established for BRSV vaccines. Some groups studying CSF virus have suggested that glycosylation is essential to maintain the integrity of neutralising epitopes, can enhance the immune response or attenuate virulent strains; beneficial criteria for vaccine antigens (27, 28). In direct contrast, vaccination with a glycosylated version of BRSV G was seen to augment disease severity (29) and glycosylation is speculated to curtail immune detection (4).

In contrast to the BRSV G protein, the F protein is highly conserved (30). It is synthesized as an inactive uncleaved precursor of 574 amino acids, denoted F0. Proteolytic cleavage at basic residues 106-109 (KKRKRR) and 131-136 (RARR) results in the production of two subunits, F1 and F2 at the carboxyl and amino termini, respectively. The F protein is an attractive vaccine antigen as it is responsible for viral entry, subsequent cell to cell viral transfer and can act as the attachment protein in the absence of G (31). The primary structure of the BRSV F protein can be observed in Appendix 5.

Peptide F_295-309_ identified in this study was observed as being reactive with BRSV antisera and is located adjacent to a known highly conserved neutralising epitope spanning F_164-293_ (32). Peptide F_295-309_ also corresponds with a region of the HRSV A2 strain F protein which was found to contain linear epitopes(33). Early studies using antisera raised in guinea pigs against a peptide spanning F_216-236_ reported efficient HRSV neutralisation *in vitro* (34), whilst Bourgeois *et al* demonstrated that residues F_200-225_ and F_255-278_ were targets for virus neutralising activity using a series of monoclonal antibodies (MAbs) (35). Additional studies identified several antigenic epitopes on the primary structure of HRSV F by analysing the binding of human polyclonal antisera to synthetic peptides (36) and van Bleek *et al* observed that human CD4+ T-cell epitopes were distributed throughout the HRSV F protein (37). However, in contrast, BRSV F-specific CD4+ T-cell epitopes were shown to be primarily located within the F1 subunit (38), despite the considerable genetic similarities in both RSV F proteins. This proposal on the location of T-cell epitopes is aligned with the data provided herein for B-cell epitopes, as the reactive peptide identified in this study is also located within the F1 subunit of BRSV. This suggests that the F1 subunit could induce T and B-cell responses in parallel and should be considered further in the development of an efficacious BRSV peptide vaccine.

It is further established that although RSV B and T-cell epitopes can be discrete, they may also be found within the same sequence. Using MAbs directed at F1 subunit-based peptides, Bouregois *et al* identified 2 antigenic domains on the HRSV F protein. Two T-cell epitopes were discovered adjacent to these neutralising epitopes (39) while peptides covering F_200-225_ and F_255-278_ were defined as targets for virus neutralising activity (36). Subsequent immunisation with these two peptides stimulated a proliferative T-cell response in BALB/c mice (40). Although peptides spanning these residues were not selected by antigenicity screening during this study, the knowledge that overlapping B and T-cell epitopes exist for RSV implies that future development of a sub-unit vaccine comprising one or two peptides capable of eliciting a broader spectrum of immunity may be possible given further research.

Phylogenetic analysis based on the G gene (Figure 1) revealed that the BRSV strain used in the present study displayed similarity to BRSV strains circulating in England in the mid-1970s but also to strains found in Missouri, Iowa and North Carolina in the late 1970s/mid 1980s, highlighting the potential use peptides identified herein may be in a global vaccine, not just one targeting the European market. Further analysis of many more American and European strains could identify this. However, phylogenetic analysis based on the F gene sequence (Figure 2) indicated that this sequence was more geographically limited to European strains of BRSV only, with a close relationship being observed between early Belgian and Dutch strains. Interestingly only a minor relationship was noted between our BRSV strain and European strains from the same time.

Antigenic heterogeneity is a common reason for vaccine failure and a major obstacle to efficacious vaccine design (41-43). A relatively broad homology across all analysed strains was demonstrated after sequence analysis of the 4 peptides identified within this study as all displayed a minimum of 71 % identity. As it is essential to any vaccine longevity that the antigen is relatively conserved peptides G_234-250_ and F_295-250_ are particularly attractive as they have a high identity to all other strains tested (>80 % identity in 100 % of strains tested).

However, many predicted peptides from this study were not antigenic when tested *in vitro* and this is likely due to a combination of the shortcomings of prediction software and the conformational nature of most B-cell epitopes, especially in the BRSV F protein. Thus, the lack of reactivity with BRSV seropositive sera against 17 peptides is not unexpected.

B-cell epitopes can be continuous, where the antigenic sequence is found in a linear fashion, or conformational (discontinuous) where the antigenic residues are distributed throughout the domain and are only brought into proximity to each other due to protein folding. A recent study into antigen-antibody binding for 53 B-cell epitopes of Influenza A and HIV obtained from Protein Data Bank, observed that none were found to be linear (44) and data suggest that the B-cell epitopes of RSV F are also mainly conformational in nature (45), relying upon the conformational structure to maintain their antigenic integrity. Arbiza *et al* observed that several epitopes of HRSV F, which had been defined as antigenic through the use of escape mutants, were not recognised by the same antibody when the epitope was synthesized as a linear peptide (46). To support this, another study described using naturally infected human sera that contained virus neutralising antibodies raised against the native conformation of the HRSV F protein. In that study the sequence of known antigenic site II on the HRSV F protein – the epitope targeted by palivizumab - was synthesized as a series of overlapping linear peptides, yet only one bound to the antibodies in the sera when this epitope was linearized (36). More recently, Jaberolansar *et al* provided evidence that linear peptides were in fact antigenic and reactive with the monoclonal antibodies used in their investigations but that the antibodies generated *in vivo* were not able to neutralise virus (47).

The online antigenicity prediction tools used here unexpectedly suggested several peptides as potentially antigenic which were highly unlikely to be so, namely, those included in the cytoplasmic/transmembrane regions of BRSV G, or those around the cleavage site or incorporating the cleavage peptide of the BRSV F protein. Similar to many other prediction algorithms, the software used herein looked at hydrophobicity/hydrophilicity of the amino acid sequence presented but it does not evaluate the structural location of the residues within the respective protein. Thus, although the algorithms work on the basis of individual amino acid properties to predict antigenic peptides, it is only from having a detailed knowledge of the tertiary protein structure could it be deduced whether these peptides were likely to be antigenic. Furthermore, a high level of non-specific binding was evident for several peptides, as evidenced by the reactivity detected regardless of whether the sera used were positive or negative for anti-BRSV antibodies. Several theories were assessed in an attempt to explain this non-specific binding, including the use of streptavidin-coated plates (48) or polyclonal instead of monoclonal sera (49) but ultimately it was not possible to rationalise why.

While currently the inherent properties within the biochemical components of a primary amino acid sequence can be used to predict the antigenicity, its immunogenicity needs to be proven. This relies on a complex, often poorly understood interaction between the peptide and the immune system (50). Often peptides can demonstrate antigenicity in vitro but this does not translate well to immunogenicity *in vivo*. Invariably, peptides require a carrier protein conjugate and an adjuvant in their vaccine formulation to induce an immune response and their necessary inclusion is an important consideration for the cost of livestock vaccine development.

Interestingly, data in the literature indicate that although many candidate peptide vaccines fail to elicit *in vitro*-detectable virus neutralising antibodies, this lack of VN activity is not always reflective of the *in vivo* protective efficacy. When calves were immunized with a previously identified antigenic peptide located within the central conserved domain of the BRSV G protein (G_174-187_) the antibodies generated lacked any virus neutralising (VN) activities *in vitro*. However, less pneumonic lung lesions were observed at post-mortem and reduced viral titres were noted in the lung washings of 2/4 calves in the vaccinated group compared to none in the control group (51). Other groups have reported protection from HRSV challenge in mice after intraperitoneal passive transfer of anti-G peptide antibodies despite no observation of virus neutralising ability *in vitro* (24, 52).

In conclusion, peptide vaccines have been reported as being less likely to induce allergic or autoimmune reactions due to the lack of redundant virus components (53) and studies performed *in vitro*, such as the one detailed here, have provided invaluable information on the specific disease-causing and protective epitopes of the virus. Although the immunogenicity of the BRSV epitopes defined herein, and their associated protective efficacy in a vaccine is yet to be defined, they provide the rationale and a solid basis for future investigation into a safer, more efficacious, sub-unit BRSV vaccine.

---

**Appendix 1:**
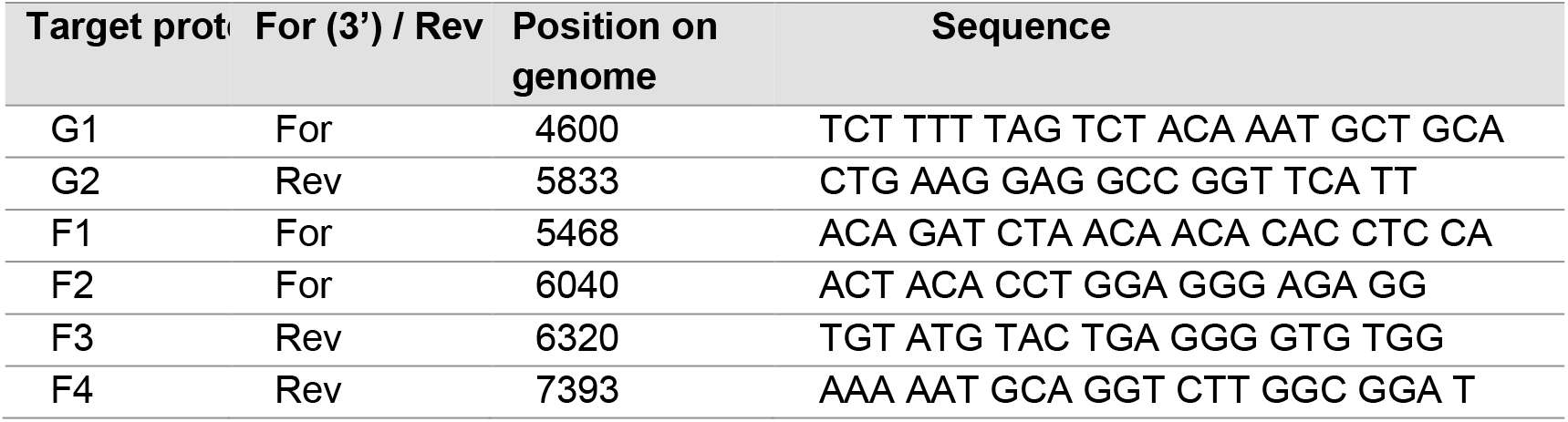
Sequence details of primers used to amplify BRSV G and F genes

**Appendix 2:**
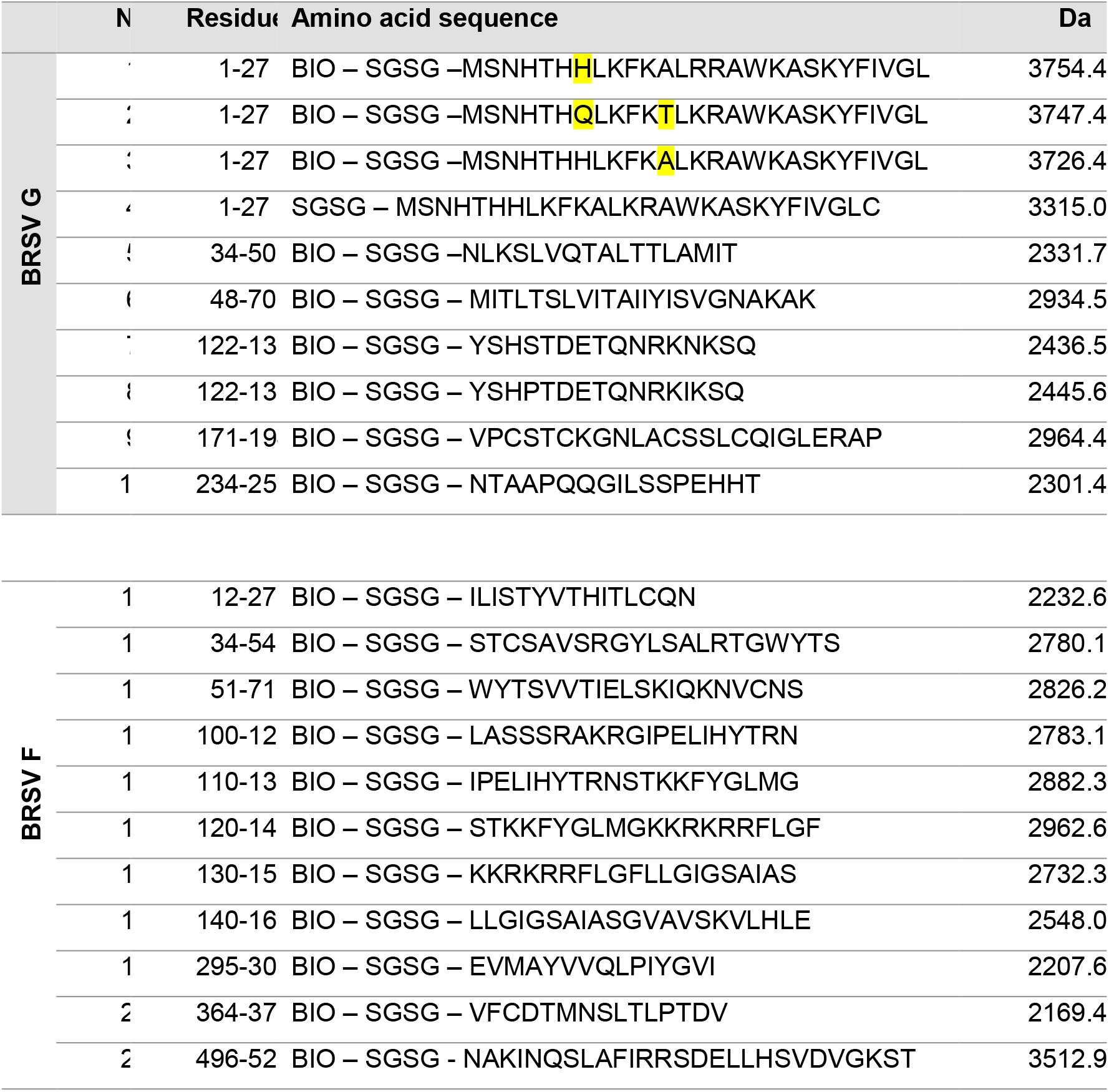
Peptides selected from BRSV G and F proteins for antigenicity screening. Highlighted text represents substitutions in amino acid residue between similar peptides. Da is the theoretical molecular weight in Daltons (Da). BIO prior to a sequence indicates a biotin tag at the N terminus.

*Appendix 3: ELISA protocol*

1. Peptides were coated 5 μg/mL in PBS 1X pH 7.4 and left to incubate at +4°C overnight. The following morning liquid was removed and the plates washed 5 times with 300 μL/well washing buffer. The final wash was left in the plate for 1 minute before discarding.
2. Two hundred μL/well of 4 % (w/v) milk solid, prepared in PBS 1X pH 7.4, was added as a blocking buffer.
3. Plates were placed in an incubator set at 37 °C for 1 h after which time the excess blocking buffer was shaken off and plates were washed as before.
4. Dilutions of bovine sera and conjugated secondary antibody were prepared in PBS 1X pH 7.4. Fifty μL/well of a 1:200 bovine sera dilution was added and plates were incubated at 37 °C for 1 h, after which time the excess serum was shaken off and plates were washed. Fifty μL/well of rabbit anti-bovine IgG (Sigma, UK) was added, used at a dilution of 1:20,000 and plates were incubated at 37 °C for 1 h.
5. For a final time, the excess liquid was shaken off and plates were washed as previously described, before the addition of 50 μL/well of TMB substrate (Biopanda Diagnostics, UK).
6. Plates were sealed and left to incubate at room temperature (RT) in the dark.
7. After 15 min, 50 μL/well 1 M H3PO4 was added to stop the reaction. Optical density (OD) was read at 450 nm using a Sunrise plate reader with Magellan software (Tecan, Switzerland).

**Appendix 4:**
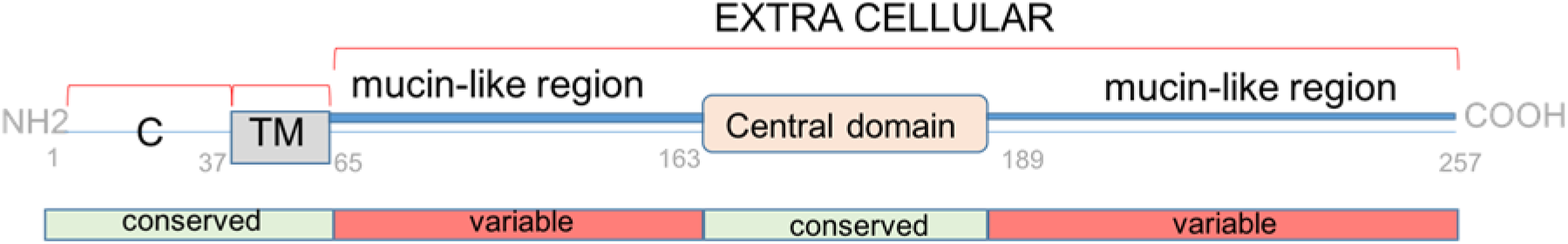
Schematic diagram of BRSV G protein primary structure C denotes the cytoplasmic domain and TM the transmembrane domain. Variable regions, corresponding to mucin-like regions are highlighted in red and amino acid numbers to delimit each domain are noted.

**Appendix 5:**
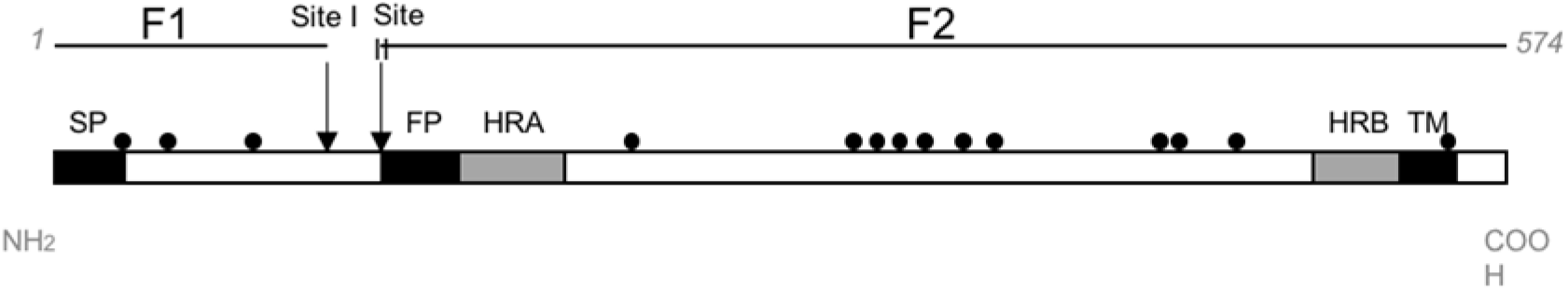
Schematic diagram of the primary structure of BRSV F protein. Black areas indicate hydrophobic domains. Also included are heptad repeat regions (HRA/HRB) and cysteine residues (black circles). The signal peptide (SP) fusion peptide (FP) and transmembrane domain (TM) are noted. Modified from (54).

## Notes

### Competing Interest Statement

The authors have declared no competing interest.

## References

1. Gershwin LJ, Schelegle ES, Gunther RA, Anderson ML, Woolums AR, Larochelle DR, et al. A bovine model of vaccine enhanced respiratory syncytial virus pathophysiology. Vaccine. 1998;16(11-12):1225–36.

2. Langedijk JP, de Groot BL, Berendsen HJ, van Oirschot JT. Structural homology of the central conserved region of the attachment protein G of respiratory syncytial virus with the fourth subdomain of 55-kDa tumor necrosis factor receptor. Virology. 1998;243(2):293–302.

3. Bukreyev A, Yang L, Fricke J, Cheng L, Ward JM, Murphy BR, et al. The secreted form of respiratory syncytial virus G glycoprotein helps the virus evade antibody-mediated restriction of replication by acting as an antigen decoy and through effects on Fc receptor-bearing leukocytes. Journal of virology. 2008;82(24):12191–204.

4. Guzman E, Taylor G. Immunology of bovine respiratory syncytial virus in calves. Mol Immunol. 2015;66(1):48–56.

5. Schlender J, Zimmer G, Herrler G, Conzelmann KK. Respiratory syncytial virus (RSV) fusion protein subunit F2, not attachment protein G, determines the specificity of RSV infection. J Virol. 2003;77(8):4609–16.

6. Taylor G, Thomas LH, Furze J, Cook RS, Wyld S, Lerch R. Recombinant vaccinia viruses expressing the F, G or N, but not M2, protein of bovine respiratory syncytial virus (BRSV) induce resistance to BRSV challenge in the calf and protect against the development of pneumonic lesions. Journal of General Virology. 1997;78(12):3195–206.

7. Sparer TE, Matthews S, Hussell T, Rae AJ, Garcia-Barreno B, Melero JA, et al. Eliminating a Region of Respiratory Syncytial Virus Attachment Protein Allows Induction of Protective Immunity without Vaccine-enhanced Lung Eosinophilia. Journal of Experimental Medicine. 1998;187(11):1921–6.

8. Langeveld JP, Casal JI, Osterhaus AD, Cortés E, de Swart R, Vela C, et al. First peptide vaccine providing protection against viral infection in the target animal: studies of canine parvovirus in dogs. Journal of virology. 1994;68(7):4506–13.

9. Greenwood DLV, Dynon K, Kalkanidis M, Xiang S, Plebanski M, Scheerlinck J-PY. Vaccination against foot-and-mouth disease virus using peptides conjugated to nano-beads. Vaccine. 2008;26(22):2706–13.

10. Dong X-N, Wei K, Liu Z-Q, Chen Y-H. Candidate peptide vaccine induced protection against classical swine fever virus. Vaccine. 2002;21:167–73.

11. Campo MS, O’Neill BW, Grindlay J, Curtis F, Knowles G, Chandrachud L. A Peptide Encoding a B-Cell Epitope from the N-Terminus of the Capsid Protein L2 of Bovine Papillomavirus-4 Prevents Disease. Virology. 1997;234:261–6.

12. Robson B. COVID-19 Coronavirus spike protein analysis for synthetic vaccines, a peptidomimetic antagonist, and therapeutic drugs, and analysis of a proposed achilles’ heel conserved region to minimize probability of escape mutations and drug resistance. Comput Biol Med. 2020;121:103749.

13. Lin L, Ting S, Yufei H, Wendong L, Yubo F, Jing Z. Epitope-based peptide vaccines predicted against novel coronavirus disease caused by SARS-CoV-2. Virus Res. 2020;288:198082.

14. Valarcher JF, Schelcher F, Bourhy H. Evolution of bovine respiratory syncytial virus. Journal of virology. 2000;74(22):10714–28.

15. van Oirschot JT. Diva vaccines that reduce virus transmission. Journal of Biotechnology. 1999;73(2):195–205.

16. Uttenthal A, Parida S, Rasmussen TB, Paton DJ, Haas B, Dundon WG. Strategies for differentiating infection in vaccinated animals (DIVA) for foot-and-mouth disease, classical swine fever and avian influenza. Expert Review of Vaccines. 2014;9(1):73–87.

17. Langedijk JP, Meleon RH, Taylor G, Furze JM, Van Oirschot JT. Structure of the Central Conserved Region of Protein G of Bovine Respiratory Syncytial Virus. Journal of Virology. 1997(May):4055–61.

18. Battles MB, McLellan JS. Respiratory syncytial virus entry and how to block it. Nat Rev Microbiol. 2019;17(4):233–45.

19. Furze J, Wertz G, Lerch R, Taylor G. Antigenic heterogeneity of the attachment protein of bovine respiratory syncytial virus. J Gen Virol. 1994;75 (Pt 2):363–70.

20. Prozzi D, Walravens K, Langedijk JP, Daus F, Kramps JA, Letesson JJ. Antigenic and molecular analyses of the variability of bovine respiratory syncytial virus G glycoprotein. J Gen Virol. 1997;78 (Pt 2):359–66.

21. Trudel M, Nadon F, Seguin C, Binz I. Protection of BALB/c mice from respiratory syncytial virus infection by immunization with a synthetic peptide derived from the G glycoprotein. Virology. 1991;165:749–57.

22. Power UF, Plotnicky-Gilquin H, Huss T, Robert A, Trudel M, Ståhl S, et al. Induction of protective immunity in rodents by vaccination with a prokaryotically expressed recombinant fusion protein containing a respiratory syncytial virus G protein fragment. Virology. 1997;230(2):155–66.

23. Nguyen TN, Power UF, Robert A, Haeuw J-F, Helffer K, Perez A, et al. The Respiratory Syncytial Virus G Protein Conserved Domain Induces a Persistent and Protective Antibody Response in Rodents. PLoS ONE. 2012;7(3):1–11.

24. Plotnicky-Gilquin H, Goetsch L, Huss T, Champion T, Beck A, Haeuw JF, et al. Identification of multiple protective epitopes (protectopes) in the central conserved domain of a prototype human respiratory syncytial virus G protein. Journal of virology. 1999;73(7):5637–45.

25. Siegrist CA, Plotnicky-Gilquin H, Cordova M, Berney M, Bonnefoy JY, Nguyen TN, et al. Protective efficacy against respiratory syncytial virus following murine neonatal immunization with BBG2Na vaccine: Influence of adjuvants and maternal antibodies. JOURNAL OF INFECTIOUS DISEASES. 1999;179(6):1326–33.

26. McGill JL, Kelly SM, Kumar P, Speckhart S, Haughney SL, Henningson J, et al. Efficacy of mucosal polyanhydride nanovaccine against respiratory syncytial virus infection in the neonatal calf. Scientific Reports. 2018;8(1):1–15.

27. Gavrilov BK, Rogers K, Fernandez-Sainz IJ, Holinka LG, Borca MV, Risatti GR. Effects of glycosylation on antigenicity and immunogenicity of classical swine fever virus envelope proteins. Virology. 2011;420(2):135–45.

28. Fernandez-Sainz I, Holinka LG, Gavrilov BK, Prarat MV, Gladue D, Lu Z, et al. Alteration of the N-linked glycosylation condition in E1 glycoprotein of Classical Swine Fever Virus strain Brescia alters virulence in swine. Virology. 2009;386(1):210–6.

29. Fuentes S, Coyle EM, Golding H, Khurana S. Nonglycosylated G-Protein Vaccine Protects against Homologous and Heterologous Respiratory Syncytial Virus (RSV) Challenge, while Glycosylated G Enhances RSV Lung Pathology and Cytokine Levels. J Virol. 2015;89(16):8193–205.

30. Stine LC, Hoppe DK, Kelling CL. Sequence conservation in the attachment glycoprotein and antigenic diversity among bovine respiratory syncytial virus isolates. Veterinary microbiology. 1997;54(3-4):201-21.

31. Karron RA, Buonaguiro DA, Georgiu AF, Whitehead SS, Adamus JE, Clements-Mann ML. Respiratory syncytial virus (RSV) SH and G proteins are not essential for viral replication in vitro: clinical evaluationv and molecular characteristics of a cold-passaged, attenuated RSV subgroup B mutant Proc Natl Acad Sci USA. 1997;94(25):13961–6.

32. López JA, Peñas C, García-Barreno B, Melero JA, Portela A. Location of a highly conserved neutralizing epitope in the F glycoprotein of human respiratory syncytial virus. Journal of virology. 1990;64(2):927–30.

33. Collins PL, E.J. CJ. Respiratory syncytial virus and metapneumovirus. In: D. M. Knipe DM, Howley PM, Griffin DE, Lamb RA, Martin MA, Roizman B, et al., editors. Field’s Virolog. 5th Ed. Philadelphia, USA: Lippincott Williams and Wilkins; 2007. p. 1601–46.

34. Trudel M, Nadon F, Seguin C, Dionne G, Lacroix M. Identification of a synthetic peptide as part of a major neutralization epitope of respiratory syncytial virus. The Journal of general virology. 1987;68 (Pt 9):2273–80.

35. Bourgeois C, Corvaisier C, Bour JB, Kohli E, Pothier P. Use of synthetic peptides to locate neutralizing antigenic domains on the fusion protein of respiratory syncytial virus. The Journal of general virology. 1991;72 (Pt 5):1051–8.

36. Scopes GE, Watt PJ, Lambden PR. Identification of a linear epitope on the fusion glycoprotein of respiratory syncytial virus. The Journal of general virology. 1990;71 (Pt 1):53–9.

37. van Bleek GM, Poelen MC, van der Most R, Brugghe HF, Timmermans HAM, Boog CJ, et al. Identification of immunodominant epitopes derived from the respiratory syncytial virus fusion protein that are recognized by human CD4 T cells. Journal of virology. 2003;77(2):980–8.

38. Fogg MH, Parsons KR, Thomas LH, Taylor G. Identification of CD4+ T cell epitopes on the fusion (F) and attachment (G) proteins of bovine respiratory syncytial virus (BRSV). Vaccine. 2001;19(23-24):3226–40.

39. Corvaisier C, Guillemin G, Bourgeois JB, Bour JB, Kohli E, Pothier P. Identification of T-cell epitopes adjacent to neutralizing antigenic domains on the fusion protein of respiratory syncytial virus. Research in Virology. 1993;144:141–50.

40. Corvaisier C, Bourgeois C, Pothier P. Cross-reactive and group-specific immune responses to a neutralizing epitope of the human respiratory syncytial virus fusion protein. Archives of Virology. 1997;142(6):1073–86.

41. Alam SM, Amin R, Rahman MZ, Hossain MA, Sultana M. Antigenic heterogeneity of capsid protein VP1 in foot-and-mouth disease virus (FMDV) serotype Asia 1. Adv Appl Bioinform Chem. 2013;6:37–46.

42. Dutta SK, Vemulapalli R, Biswas B. Deficiency in antibody response to vaccine and heterogeneity of Ehrlichia risticii strains with Potomac Horse Fever vaccine failure in horses. Journal of Clinical Microbiology. 1997;Feb 1998:506–12

43. Kumagai A, Kawauchi K, Andoh K, Hatama S. Sequence and unique phylogeny of G genes of bovine respiratory syncytial viruses circulating in Japan. Journal of Veterinary Diagnostic Investigation. 2021;33(1):162–6.

44. Rubinstein ND, Mayrose I, Halperin D, Yekutieli D, Gershoni JM, Pupko T. Computational characterization of B-cell epitopes. Molecular Immunology. 2008;45(12):3477–89.

45. Davidson E, Doranz BJ. A high-throughput shotgun mutagenesis approach to mapping B-cell antibody epitopes. Immunology. 2014;143(1):13–20.

46. Arbiza J, Taylor G, López JA, Furze J, Wyld S, Whyte P, et al. Characterization of two antigenic sites recognized by neutralizing monoclonal antibodies directed against the fusion glycoprotein of human respiratory syncytial virus. The Journal of general virology. 1992;73 (Pt 9):2225–34.

47. Jaberolansar N, Chappell KJ., Watterson D, Bermingham IM, Toth I, Young PR, et al. Induction of high titred, non-neutralising antibodies by self-adjuvanting peptide epitopes derived from the respiratory syncytial virus fusion protein. Scientific Reports. 2017;7(1):1–11.

48. Välimaa L, Laurikainen K. Comparison study of streptavidin-coated microtitration plates. Journal of Immunological Methods. 2006;308(1):203–15.

49. Lipman NS, Jackson LR, Trudel LJ, Weis-Garcia F. Monoclonal versus polyclonal antibodies: Distinguishing characteristics, applications, and information resources. ILAR JOURNAL. 2005;46(3):258–68.

50. Van Regenmortel MH. Antigenicity and immunogenicity of synthetic peptides. Biologicals : journal of the International Association of Biological Standardization. 2001;29(3-4):209–13.

51. Bastien N, Taylor G, Thomas LH, Wyld SG, Simard C, Trudel M. Immunization with a peptide derived from the G glycoprotein of bovine respiratory syncytial virus (BRSV) reduces the incidence of BRSV-associated pneumonia in the natural host. Vaccine. 1997;15(12-13):1385–90.

52. Power UF, Plotnicky-Gilquin H, Goetsch L, Champion T, Beck A, Haeuw J-F, et al. Identification and characterisation of multiple linear B cell protectopes in the respiratory syncytial virus G protein. Vaccine. 2001;19(17):2345–51.

53. Skwarczynski M, Toth I. Peptide-based synthetic vaccines. CHEMICAL SCIENCE. 2016;7(2):842–854

54. Melero JA, Mas V, McLellan JS. Structural, antigenic and immunogenic features of respiratory syncytial virus glycoproteins relevant for vaccine development. Vaccine. 2017;35(3):461–8.

